# Brainstem premotor mechanisms underlying vocal production and vocal-respiratory coordination

**DOI:** 10.1101/2023.10.12.562111

**Authors:** Jaehong Park, Seonmi Choi, Jun Takatoh, Shengli Zhao, Andrew Harrahill, Bao-Xia Han, Fan Wang

## Abstract

Speech generation critically depends on precise controls of laryngeal muscles and coordination with ongoing respiratory activity. However, the neural mechanisms governing these processes remain unknown. Here, we mapped laryngeal premotor circuitry in adult mice and viral-genetically identified excitatory vocal premotor neurons located in the retroambiguus nucleus (RAm^VOC^) as both necessary and sufficient for driving vocal-cord closure and eliciting mouse ultrasonic vocalizations (USVs). The duration of RAm^VOC^ activation determines the lengths of USV syllables and post-inspiration phases. RAm^VOC^-neurons receive inhibitory inputs from the preBötzinger complex, and inspiration needs can override RAm^VOC^-mediated vocal-cord closure. Ablating inhibitory synapses in RAm^VOC^-neurons compromised this inspiration gating of laryngeal adduction, resulting in de-coupling of vocalization and respiration. Our study revealed the hitherto unknown circuits for vocal pattern generation and vocal-respiratory coupling.

**One-Sentence Summary:** Identification of RAm^VOC^ neurons as the critical node for vocal pattern generation and vocal-respiratory coupling.

## Main Text

Vocalization is used in many species for social communications. In mammals, vocal sounds are produced by the air blowing out from the lungs against the near-closed vocal cords in the larynx (*1*). Hence, vocalization needs to be coordinated with breathing: the larynx is only closed (adduction of the vocal cords) during expiration for producing sound, while inspiration opens the larynx to inflate the lungs (to bring in enough air) for the next bout of vocalization. Premature closing or opening of the vocal cords can lead to inspiration problems and coarse vocalization (*2, 3*), respectively. While previous studies have uncovered higher centers regulating vocalization, such as the gating of vocal production by neurons in the periaqueductal gray (PAG) (*4, 5*); and the modulation of vocalization by social-cues (*6, 7*) and group interactions (*8, 9*) by neurons in the hypothalamus and motor cortex in different species, the precise neural mechanisms for producing the actual sounds via vocal cord adduction and vocal-breathing coordination remain unknown.

The movement of the vocal cords is controlled by laryngeal motoneurons, whose activity patterns are generated by premotor circuits in the brainstem (*10, 11*). A recent study revealed an intermediate reticular oscillator (iRO) in neonatal mice that synchronizes neonatal crying with breathing (*12*), although whether iRO is involved in adult vocalization remains unclear. Many other studies that used electrical, pharmacological, or electrophysiology methods suggested that the caudal-ventral brainstem area, specifically the <u>r</u>etro<u>am</u>biguus nucleus (RAm), is a key node in vocal pattern generations in adults (*13–16*). The RAm region does not have anatomical demarcations, and likely contains heterogeneous types of neurons. Thus, past studies did not identify vocal-specific premotor neurons/circuits within the RAm, and consequently non-specific manipulations of RAm did not result in naturalistic vocalizations (*14, 16, 17*). Regarding the vocal-breathing coordination, while intensive research has been conducted on breathing controls (*18–20*) ever since the discovery of the inspiration rhythm generator, the preBötzinger complex (preBötC) (*21*), decades ago, very few studies investigated the function of the preBötC during vocalizations in awake animals (*22*). Therefore, how preBötC interacts with laryngeal motoneurons and/or premotor neurons to coordinate breathing and vocalization is unknown. Here, we identified vocal-specific laryngeal adduction premotor neurons in RAm (RAm^VOC^) through transsynaptic tracing, activity-dependent labeling, and functional manipulations, along with behavioral studies and laryngeal imaging in adult mice. We further investigated how inspiration controls the activity of RAm^VOC^ neurons for vocal-breathing coordination.

### Vocalization-specific laryngeal premotor neurons in the brainstem

The activity of laryngeal muscles and motoneurons is controlled by premotor neurons in the hindbrain (*10, 23*). However, the location and identity of the laryngeal premotor circuits in adult mammals have yet to be revealed. To this end, we applied a three-step mono-synaptic tracing strategy (*24*) (Fig. 1A-B), combining AAVretro-Cre (*25*), injected into laryngeal muscles; Cre-dependent helper AAVs to express TVA receptor (*26*) and rabies glycoprotein G (*27*) in motoneurons, and pseudotyped G-deleted rabies virus (EnvA^M21^-RV-GFP) (*28*), injected into the nucleus ambiguus (NA). Trans-synaptically labeled laryngeal premotor neurons were mostly found in the brainstem (Fig. 1C, fig. S1), specifically in the Kölliker-Fuse (KF), parvocellular reticular formation (PCRt), lateral paragigantocellular nucleus (LPGi), intermediate reticular nucleus (IRt), preBötC, nucleus tractus solitarii (NTS), and RAm (Fig. 1C). We registered all labeled neurons in the Allen common coordinate frame for the mouse brain (Allen CCF) (*29*) and compared to our previously identified maps of other orofacial muscle premotor neurons (jaw and tongue) (*24*) (fig. S1). The spatial distributions of the laryngeal premotor neurons from different mice (n=3) were similar but were distinct from those of jaw and tongue premotor maps (fig. S1). Interestingly, labeled premotor neurons also had extensive collateral projections to the other motor nuclei, including the trigeminal (5N), the facial (7N), and the hypoglossal (12N) nuclei (fig. S1), suggesting that laryngeal premotor neurons simultaneously recruit other orofacial motoneurons for vocalization and perhaps other movements.

**Fig. 1.**
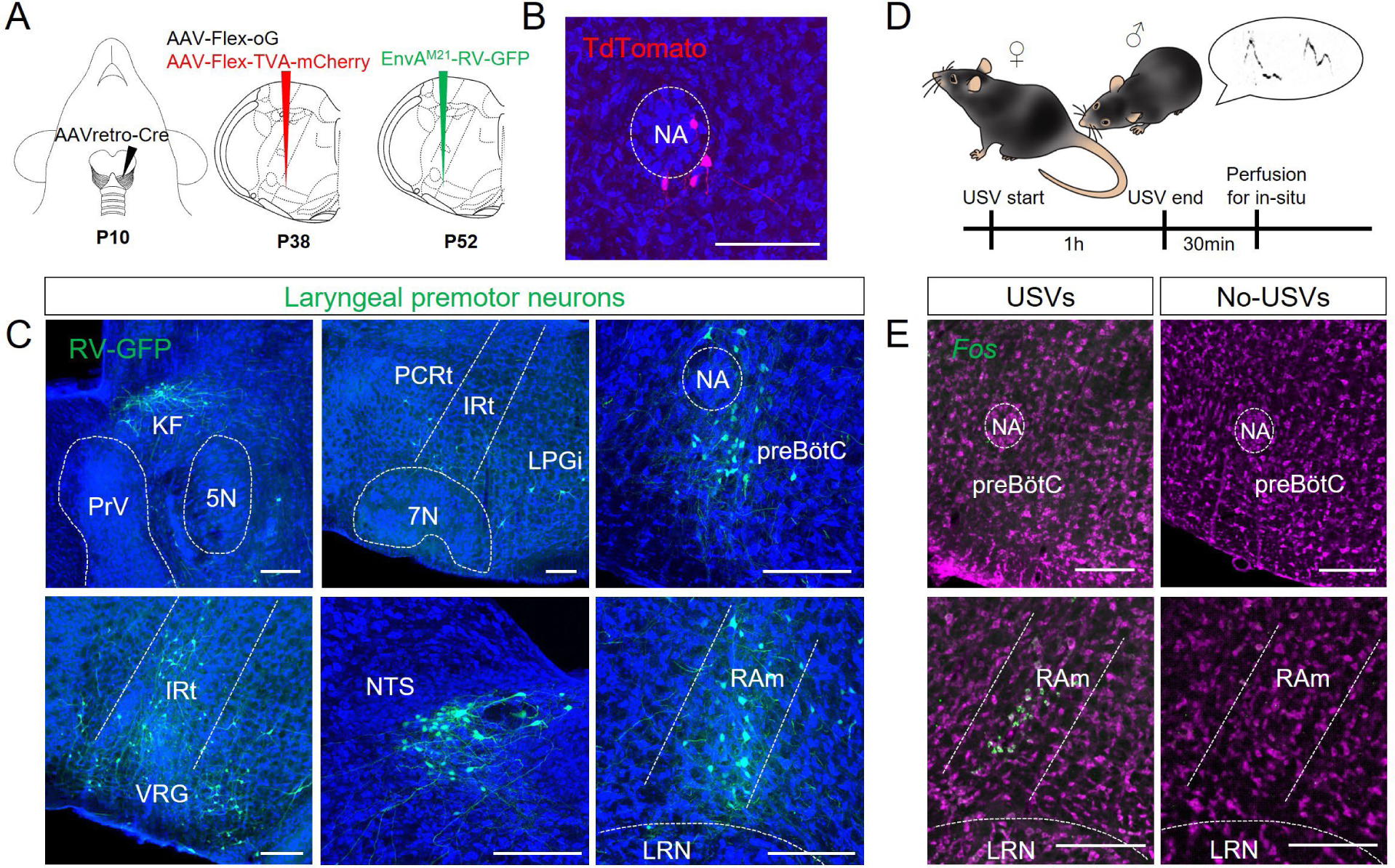
Transsynaptic mapping of laryngeal premotor neurons and vocalization-induced *Fos* activity in RAm. (A) A schematic three-step viral strategy using AAVretro-Cre, helper virus (AAV-flex-oG, AAV-flex-TVA-mCherry), and monosynaptic rabies virus (EnvA^M21^ coated) to map laryngeal premotor neurons. (B) Laryngeal motoneurons (red) labeled by AAVretro-Cre in an Ai14 reporter mouse around the nucleus ambiguus (NA). (C) Laryngeal premotor neurons (green) in the KF, PCRt, LPGi, preBötC, IRt, VRG, NTS, and RAm. (D) A scheme for *Fos* induction experiments in a social-context eliciting USVs in male mice. (E) In situ hybridization with *Fos* probe (green) in the preBötC (upper left) and RAm (bottom left) after female-directed male USVs. The right columns show *Fos* activity in the same regions without vocalization behaviors (male mice in a chamber alone). Neurotrace Blue was used to visualize neuronal structures (Blue color for B and C; magenta for E). All scale bars indicate 200 µm.

Consistent with previous suggestions that RAm is a critical node for vocal production, our premotor tracing indeed labeled a cluster of neurons in the RAm (Fig. 1C). To further explore whether RAm contains neurons activated by vocalization, we examined *Fos* mRNA expression in male mice 90-min after female-induced courtship ultrasonic vocalizations (USVs) (Fig. 1D). Robust *Fos* signals were observed in the RAm (Fig. 1E). By contrast, fewer and weaker *Fos* expressions were found in other premotor brainstem areas, such as the preBötC in the same samples (Fig. 1E). These data enticed us to further investigate the precise role of RAm laryngeal premotor neurons in vocalization.

We first used the Fos-based cell targeting method called CANE (*28*) to label courtship USV-activated Fos+ neurons in RAm in male mice (hereafter referred to as RAm^VOC^ neurons) (Fig. 2A). After labeling RAm^VOC^ neurons with GFP via CANE, we re-exposed male mice to female to re-elicit USVs and Fos expression and confirmed that labeled RAm^VOC^ were Fos+ (Fig. 2B). We further registered the locations of all CANE-captured RAm^VOC^ neurons in the Allen CCF and confirmed that their positions overlapped with those of the rabies-traced laryngeal premotor neurons in this region (Fig. 2C). To rule out the possibility that some of the CANE-captured RAm^VOC^ neurons were vocal motoneurons, we examined the expression of ChAT, a molecular marker for motoneurons. None of the RAm^VOC^ neurons expressed ChAT (Fig. 2D), i.e., they are not cholinergic motoneurons. Furthermore, the axonal boutons from the labeled RAm^VOC^ neurons innervated ChAT positive neurons around the NA (Fig. 2D), consistent with them being vocal premotor neurons. Lastly, in-situ hybridization using *Vglut2* and *Vgat* probes showed that the majority of RAm^VOC^ neurons were glutamatergic neurons (*Vglut2*+: 85.05±0.06%, *Vgat*+: 12.85±0.02%, n=3 mice, Fig. 2E) suggesting that they provide excitatory inputs to laryngeal motoneurons.

**Fig. 2.**
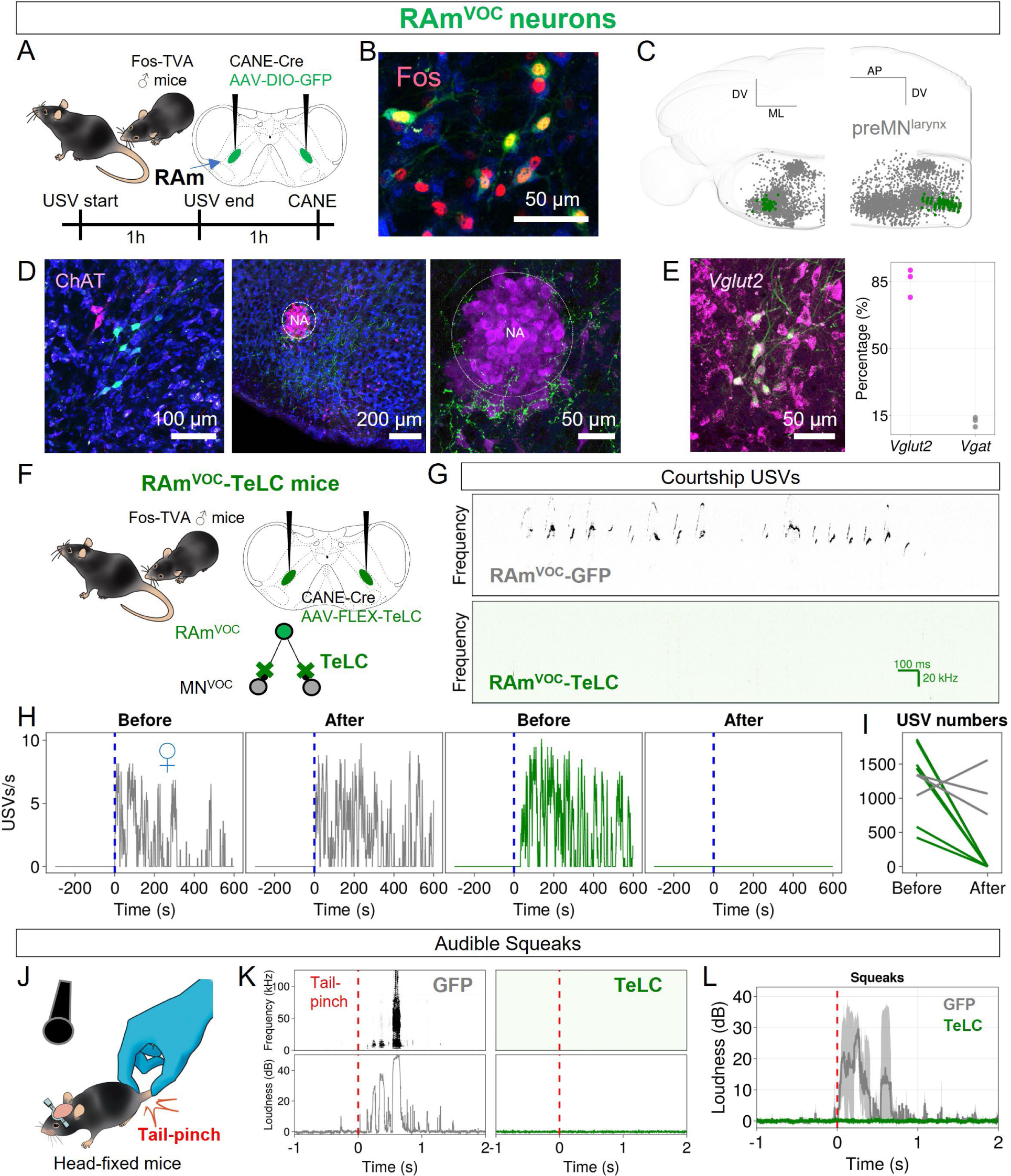
Vocalization-induced Fos positive neurons in the RAm (RAm^VOC^ neurons) are excitatory laryngeal premotor neurons and required for vocalization in mice. (A) A scheme for CANE experiments to capture vocalization-induced Fos positive neurons in the RAm. (B) RAm^VOC^ neurons (green) with Fos immunolabeling (red). (C) RAm^VOC^ neurons (green) with the laryngeal premotor neurons (grey) in the Allen CCF in coronal (left) and sagittal (right) views. (D) RAm^VOC^ neurons (green) with ChAT immunolabeling (red). Left (soma) and middle (axon terminals). The right panel highlights the NA region of the middle panel. (E) RAm^VOC^ neurons (green) with HCR in situ hybridization labeling for *Vglut2* and *Vgat* from n=3 mice. (F) A scheme for expressing TeLC in RAm^VOC^ neurons. (G) Spectrograms of female-directed USVs of RAm^VOC^-GFP controls (upper) and RAm^VOC^-TeLC (bottom) mice. (H) USV rates of male mice during courtship behaviors for 10 min. Blue vertical lines indicate the time of female introduction (♀). Grey and green plots for a representative RAm^VOC^-GFP and a RAm^VOC^-TeLC mouse, before and 2 weeks after virus injection (left and right, respectively). (I) The total numbers of USV syllables during 10min social interactions (RAm^VOC^-TeLC, green, n=6; and control, grey, n=3). (J) A scheme for recording tail pinch-induced audible squeaks. (K) Spectrogram (upper) and sound intensity plots (bottom) of RAm^VOC^-GFP and RAm^VOC^-TeLC mice. Red vertical lines indicate the onset of tail-pinch stimuli. (L) Average intensity of squeaks during tail-pinch (n=3 for each group).

### Silencing RAm^VOC^ neurons abolishes both ultrasonic and audible vocalizations

To dissect the functional role of RAm^VOC^ neurons, we bilaterally expressed tetanus toxin light chain (TeLC) to inhibit their synaptic outputs, or expressed GFP as control, in RAm^VOC^ using CANE (Fig. 2F). RAm^VOC^-GFP male mice emitted robust USVs in the presence of a female partner before and after CANE-mediated expression (Fig. 2G upper and H left). In contrast, RAm^VOC^-TeLC mice failed to vocalize in response to female mice after TeLC expression (Fig. 2G bottom and H right). Other mating behaviors, such as chasing and sniffing female mice, remained intact (data not shown). The effect of silencing RAm^VOC^ neurons was robust and consistent: all six RAm^VOC^-TeLC mice had complete mutism (zero utterance, Fig. 2I). These results show that RAm^VOC^ neurons are essential for producing USVs.

In addition to social USVs, mice also elicit audible-range squeaks in response to strongly aversive stimuli (*30*). Prior studies suggested that USVs and squeaks are triggered by different neural pathways (*5, 30*). For example, a recent study showed that inhibition of the PAG-RAm pathway only abolished USVs but not pain-elicited audible vocalizations (*5*). We evoked squeaks in mice using a tail-pinch stimulus (Fig. 2J). While control RAm^VOC^-GFP mice responded with robust squeaks, RAm^VOC^-TeLC mice were silent (Fig. 2K and L). Furthermore, when we applied foot-shocks, RAm^VOC^-GFP (Movie S1), but not RAm^VOC^-TeLC mice (Movie S2), squeaked, even though all mice exhibited escape behaviors, indicating that nociceptive responses of the RAm^VOC^-TeLC mice were intact.

To rule out the unlikely possibility that mutism in the RAm^VOC^-TeLC mice originated from breathing troubles, we habituated mice on a treadmill wheel and gently encouraged them to run (fig. S2). Running changes both the frequency and amplitude of breathing in mice (*31*). We found that the modulation of respiration by running in RAm^VOC^-TeLC mice remained intact as that in the control group (fig. S2: Amplitude changes: 28.8±14.5% (TeLC, n=3 mice) vs 24.4±5.2% (GFP, n=4 mice), p=0.8597, Mann-Whitney U test; Frequency changes: 37.7±31.5% (TeLC, n=3 mice) vs 27.3±19.1% (GFP, n=4 mice), p=0.5959, Mann-Whitney U test). Taken together, our findings indicated that RAm^VOC^ neurons are vocalization-specific but not general respiratory modulating premotor neurons, and that they are required for both audible squeaks and USVs in mice.

### RAm^VOC^ activation is sufficient to elicit and modulate species-specific vocalizations

To determine whether RAm^VOC^ neurons are sufficient to elicit species-specific vocalization, we expressed the potent optogenetic activator ChRmine (*32*) in these neurons, again using CANE in male mice (Fig. 3A). Note that vocal cord adduction is the critical step in mice vocalization, as the close distance of the vocal cords enables a jet stream of air to come through a small hole for producing USV (*33–35*). We therefore first tested the effect of optogenetic activation of RAm^VOC^ neurons on vocal cord closure. The larynx was imaged with a camera while mice were anesthetized and placed in a prone position (Fig. 3B). Measurements of the glottal area were obtained with or without optogenetic stimulation of RAm^VOC^-ChRmine neurons. The vocal cords naturally widened and narrowed (but not fully closed) rhythmically (Fig. 3C, Movie S3), in phase with inhalation and exhalation, resulting in periodic changes in the size of the glottal area (Fig. 3D). Optogenetic activation of RAm^VOC^ with 5s continuous laser illumination (560nm for ChRmine activation) instantaneously closed the vocal cords, and the laryngeal adduction persisted during the stimulation (Fig. 3D, n=3 mice, Movie S3). This prolonged laryngeal adduction was interrupted by occasional glottal openings during the 5s stimulation in all mice tested (this point is further elaborated below). We next examined the effect of RAm^VOC^ activation in awake male mice (Fig. 3E). Remarkably, applying a brief 100ms laser pulse (with 2s inter pulse intervals) reliably induced ultrasonic vocalizations time-locked to each pulse (Fig. 3F). Spectrogram analysis showed that the RAm^VOC^ activation-induced vocalizations were qualitatively similar to natural USVs (Fig. 3G). The onset latencies of the optogenetic-induced USVs were short (48.0±13.5ms, n=445 syllables from n=3 mice, Fig. 3H). We also compared the distributions of opto-evoked and spontaneous USVs for several other acoustic features, including the duration, loudness, spectral purity, mean sound frequency, and pitch variance (Fig. 3I). A principal component analysis (PCA) using those five features showed a complete overlap between opto-evoked and spontaneous USVs on the PCA space defined by the first two components (71.5% variance explained), suggesting that the RAm^VOC^ activation-evoked vocalizations closely resembled the natural USVs produced by the same mice (Fig. 3J). Taken together, these results showed that activation of RAm^VOC^ neurons is sufficient to drive vocal cord adduction and elicit USVs, and that RAm^VOC^ activation-evoked vocalizations (RAm^VOC^-USVs) were consistent with natural, species-specific USVs rather than artificial, non-specific vocalizations (that were observed in previous non-specific RAm manipulation studies (*14, 16, 17*)).

**Fig. 3.**
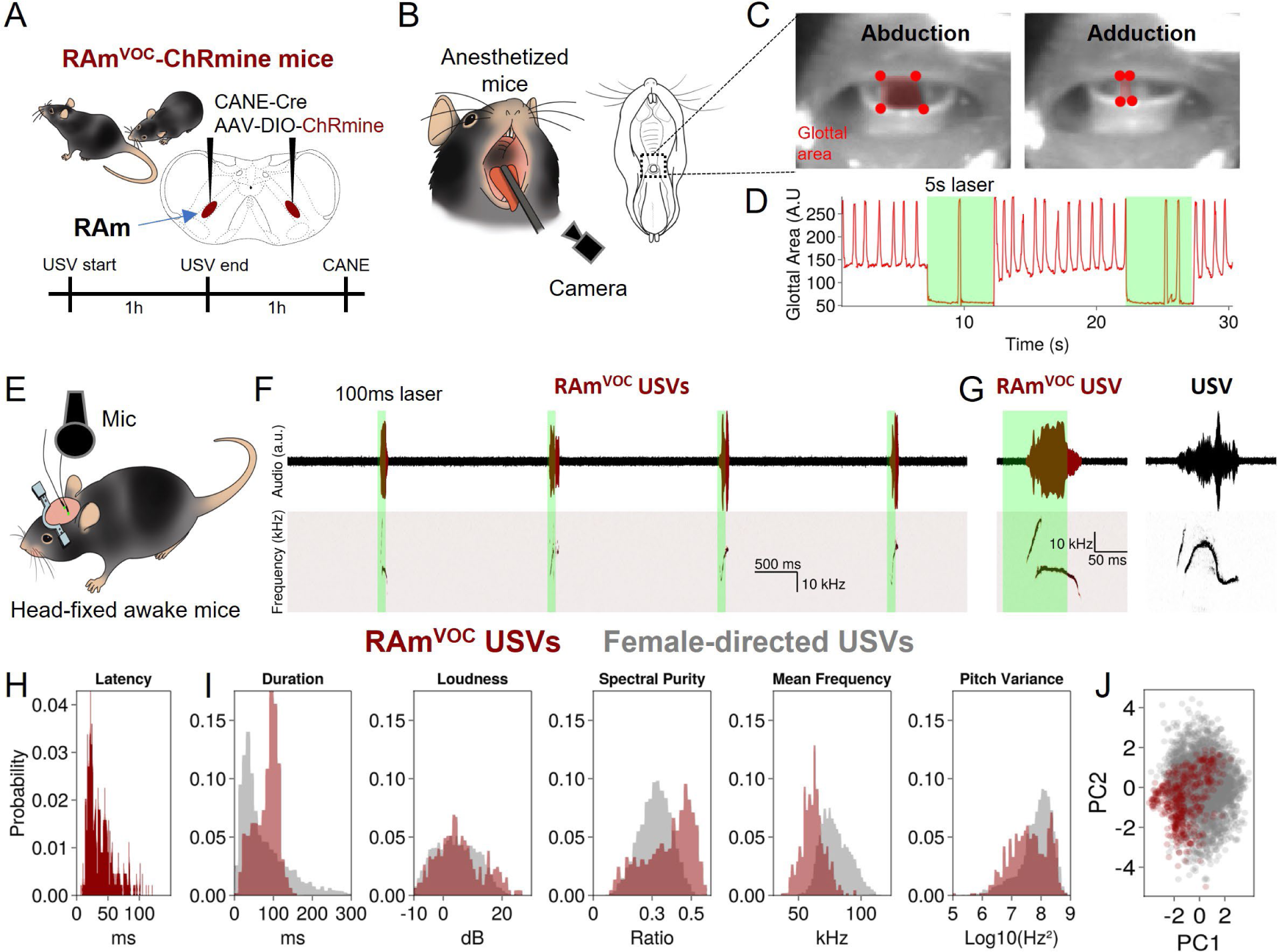
Optogenetic activation of RAm^VOC^ neurons robustly elicit species-specific USV-like vocalizations in mice. (A) Schematic for expressing ChRmine in RAm^VOC^ neurons using CANE method. (B) Schematic for visualizing vocal cords in anesthetized mice. (C) Images showing opened (left) and closed (right) vocal cords. Red dots indicate the cartilage parts of the vocal cords that are used to track the glottal area (red rectangle). (D) The response of the glottal area to RAm^VOC^ opto-activation. Green bar (5s) indicates the laser stimulation period. (E) Schematic for recording vocalization of awake mice in a head-fixed condition. (F) Sound-time raw traces (upper) and frequency-time spectrogram (bottom) during a train of brief laser pulses (laser wavelength = 560nm, 100ms of 4 pulses with 2s intervals). (G) Examples of opto-induced (left) and female-directed (right) USVs in a male RAm^VOC^ ChRmine mouse. (H) Latency distribution of opto-induced USVs (laser duration:100ms, 445 syllables from n=3 mice). (I) Distributions of five acoustic features (duration, loudness, spectral purity, mean frequency, and pitch variance) of opto-induced USV syllables (445 syllables from n=3 mice) and female-directed USVs (5411 syllables from n=3 mice). (J) A first-second PCA map of all USV syllables from five acoustic USV features.

Given that a brief RAm^VOC^ activation elicited a single short USV syllable (Fig. 3F), we also tested whether RAm^VOC^ activation can alter the patterns of USVs, specifically, the length of individual USV syllables. We varied the duration of optogenetic stimulation of RAm^VOC^ (50, 100, and 200ms), and observed that indeed the length of RAm^VOC^-USV syllables were proportionally correlated to the duration of laser stimuli (Fig. 4B and D). These data indicate that the duration of USV syllables can be controlled by the extent of the persistent RAm^VOC^ activity.

**Fig. 4.**
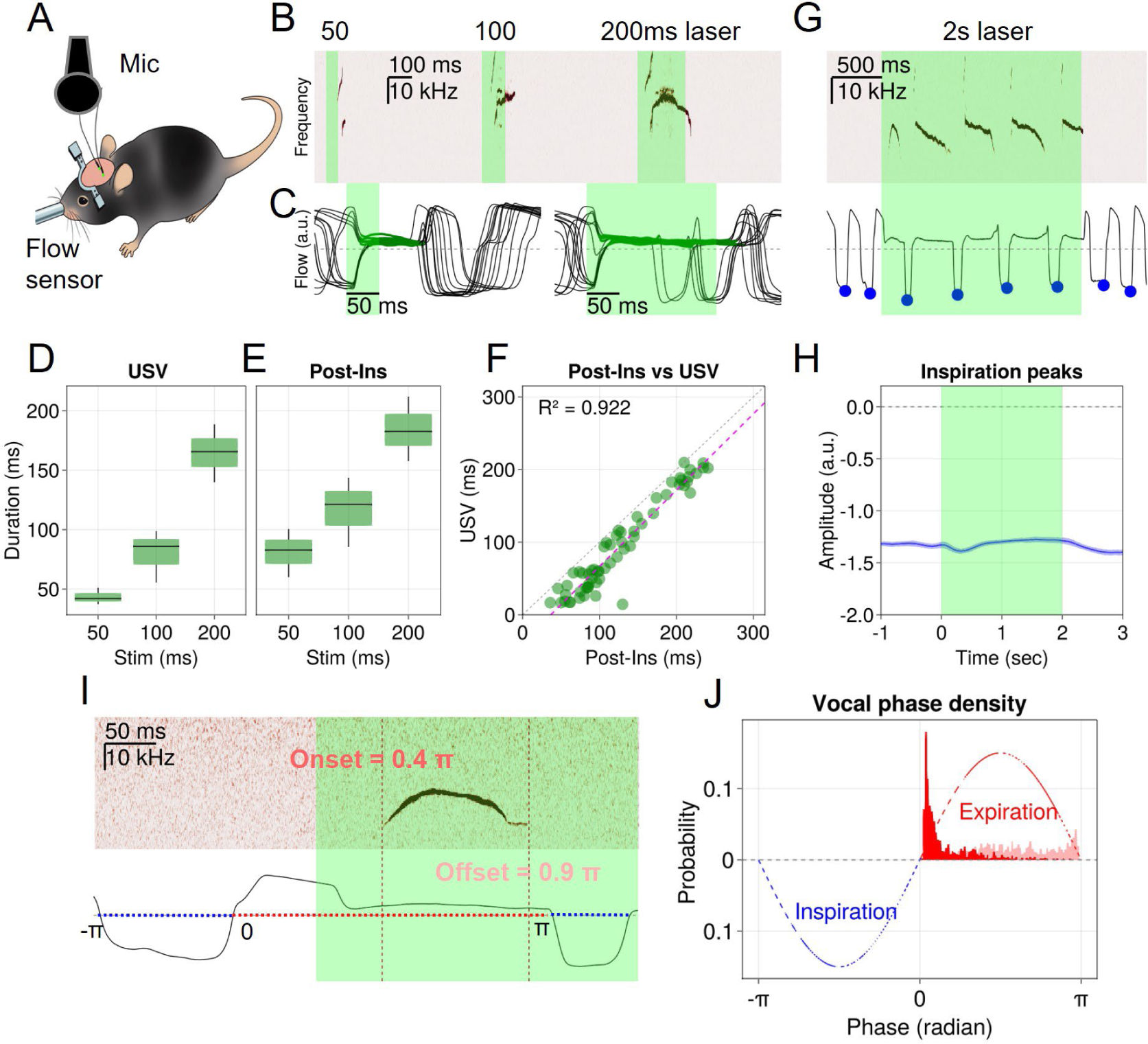
RAm^VOC^ activation can modulate the duration of USVs and post-inspiration phases (post-Ins) until interrupted by the need for breathing. (A) Schematic for recording vocalization and respiration in RAm^VOC^ ChRmine mice. (B) Representative USV syllables evoked by three different durations of RAm^VOC^ laser activation (50, 100, and 200ms). (C) Representative respiratory responses to the RAm^VOC^ activation (Left: 50ms, Right: 200ms). 13 trials are aligned to the laser onsets and overlayed. Green lines indicate the post-Ins. (D) Average duration of RAm^VOC^ USVs (n=3 mice). (E) Average duration of RAm^VOC^-induced post-Ins (n=3 mice). (F) Correlation between the duration of RAm^VOC^-induced post-Ins and USVs. A magenta dotted line indicates the linear regression of the two parameters. (G) USVs (upper) and respiratory responses (bottom) to the 2s RAm^VOC^ activation. Blue dots indicate the inspiratory flow peaks. (H) Average inspiration peak amplitudes during 2s RAm^VOC^ ChRmine stimulation. Note that discrete inspiration peaks (blue dots) were interpolated to continuous traces to calculate the mean and 95% confidence interval. (I) Projection of onset and offset of a RAm^VOC^ USV onto a respiratory phase. Inspiration: −π to 0, Expiration: 0 to π. (J) Phase density distribution of the onsets (red) and offsets (pink) of RAm^VOC^ USVs. Blue and red dash lines represent arbitrary inspiration and expiration phase, respectively.

### Vocalization-respiration interactions during RAm^VOC^ activation

For normal vocalization, sound is exclusively produced during the expiration phase, specifically during the post-inspiration phase (post-Ins). The post-Ins is the period when air flow plateaus, and vocalization extends the post-Ins with increased airway resistance by adducting the larynx to generate sound syllables (*36*). To investigate the potential role of RAm^VOC^ in vocal-respiration interaction, we simultaneously measured USVs and respiratory activity in mice while optogenetically stimulating RAm^VOC^ with different durations (50, 100, and 200ms) (Fig. 4A-C). We found that longer RAm^VOC^ activation induced longer post-Ins (Fig. 4C and E). The durations of USV syllables and post-Ins were highly correlated (R^2^=0.922, Fig. 4F), a fact consistent with the notion that RAm^VOC^ activity mediates vocal cord closure that simultaneously drives vocalization and restricts air flow.

It is also known that inspiration need – i.e., the urge to breath-in when the blood oxygen level drops - can stop vocal production. With 200ms RAm^VOC^ activation, we occasionally observed a full inspiration cycle during stimulation (200ms, Fig. 4C). Similarly, as described above, in the anesthetized larynx imaging preparation, vocal cords could be observed to occasionally open during prolonged 5s RAm^VOC^ activation, presumably due to an “override” breathing command signal (Fig. 3D). To further investigate this inspiratory interruption of vocalization/vocal adduction in awake mice, we used 2s continuous RAm^VOC^ opto-activation. This 2s stimulation produced multiple USV syllables accompanied by concurrent post-Ins, which was interrupted by intervening inspirations (Fig. 4G). The amplitudes of the intervening inspirations were similar to those in the baseline conditions, indicating that these are normal breaths (Fig. 4G and H). We projected the onsets and offsets of the multiple USV syllables evoked by the 2s RAm^VOC^ activation onto respiration phase maps (Inspiration: −π to zero, Expiration: zero to π, Fig. 4I). All syllables were exclusively found in the expiration phase (Fig. 4J), consistent with the notion that intervening inspirations stop the on-going USVs evoked by RAm^VOC^ activation.

### Inhibitory inputs to RAm^VOC^ are essential for inspiration interruption of vocalizations

We hypothesized that inhibitory inputs onto RAm^VOC^ neurons are the key for the periodic suppression of vocalization by inspiration. To identify the source of inspiration-related inhibitory inputs to the RAm^VOC^ neurons, we performed monosynaptic tracing of presynaptic neurons to RAm^VOC^ (preRAm^VOC^). This was achieved by expressing TVA and oG in RAm^VOC^ using CANE, followed by infecting these neurons with a pseudo-typed G-deleted rabies virus (Fig. 5A). Tracing results showed that RAm^VOC^ neurons receive excitatory inputs from the PAG (as expected based on prior studies), the parabrachial (PB)/Kölliker-Fuse nucleus, and other areas (Fig. 5B, and data not shown). Excitatory PAG neurons are known to be required for eliciting USVs but not for generating rhythmic vocal patterns (*5*). Interestingly, the dominant source of inhibitory inputs to RAm^VOC^ neurons was the preBötC (Fig. 5B), the inspiration rhythm generator. In our mapping of laryngeal premotor neurons, we also labeled a population of inhibitory neurons in the preBötC (fig. S3). Thus, the preBötC provides inhibitory inputs to both vocal motoneurons (MN^VOC^) and to RAm^VOC^ (Fig. 5C). These circuit tracing results suggest that the inspiration-gated periodic patterns of USVs could be generated by tonic excitatory inputs from the PAG to RAm^VOC^ to induce vocal cord adduction that is interrupted by rhythmic inhibition from the preBötC to both MN^VOC^ and RAm^VOC^ (Fig. 5C).

**Fig. 5.**
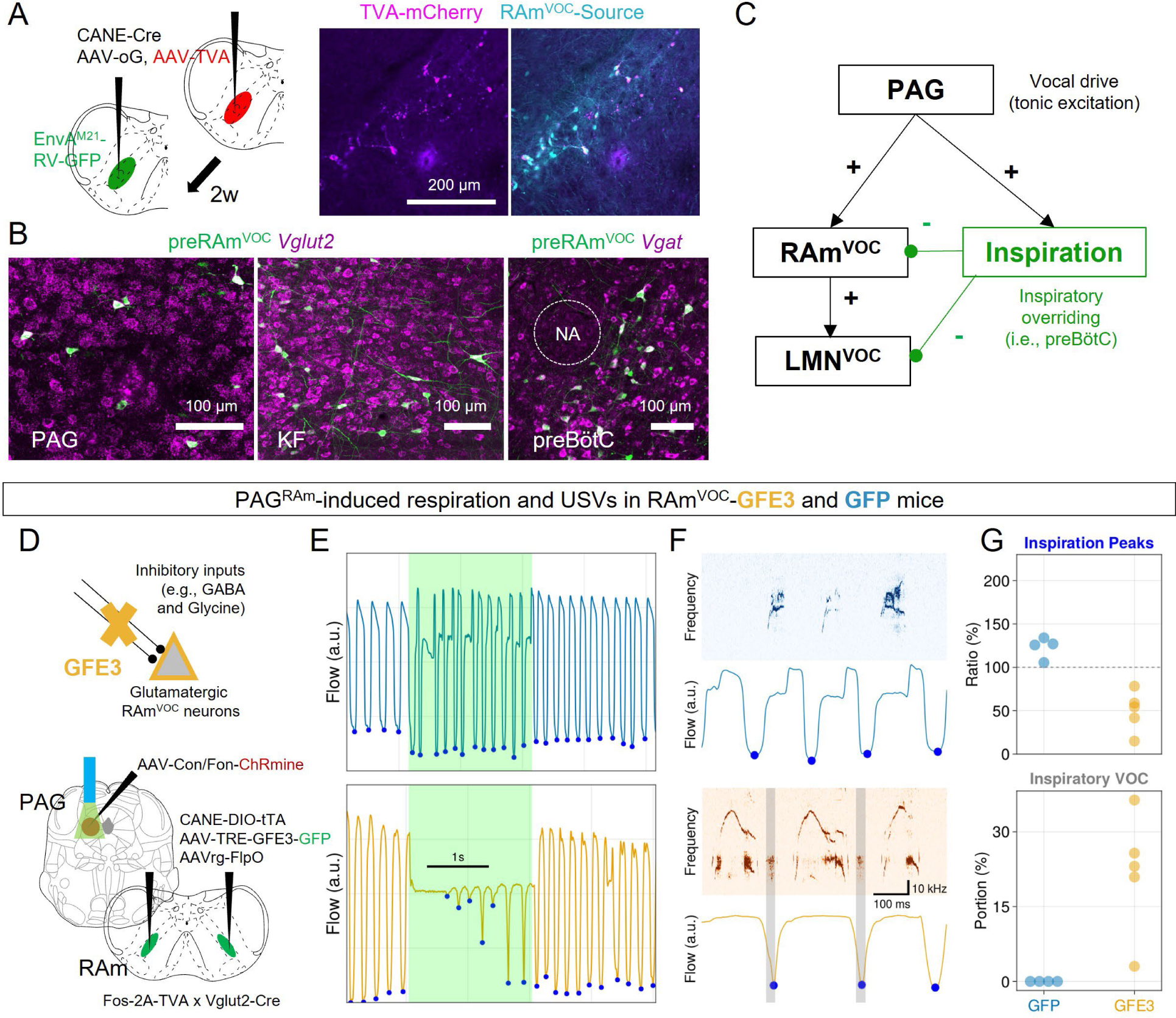
Ablating inhibitory synapses on RAm^VOC^ neurons compromised vocal-respiratory coordination. (A) Schematic for transsynaptically tracing preRAm^VOC^ neurons (left). CANE and rabies labeled source cells (magenta: TVA, cyan: GFP) in RAm (right). (B) representative images of labeled preRAm^VOC^ neurons (green) in the PAG, KF, and preBötC with in situ hybridization (magenta for *Vglut2* and *Vgat*). (C) Schematic of the proposed neural mechanism for vocal-respiratory coordination. (D) Schematic for ablating inhibitory synapses in RAm^VOC^ neurons with GFE3 expression in the glutamatergic RAm^VOC^ neurons (RAm^VOC^-GFE3), and concurrent expression of ChRmine in RAm projecting glutamatergic PAG neurons. (E) Respiratory activities of the RAm^VOC^-GFE3 and RAm^VOC^-GFP mice in response to the PAG^RAm/vglut2^-ChRmine stimulation for 2s. Blue dots represent the inspiratory peaks. (F) Spectrogram (upper) with the respiratory responses (bottom). Grey bars label abnormal vocalizations in the inspiratory phases. (G) Average changes in the inspiratory peaks of the mice (n=5, GFE3; n=4, GFP, upper) during the PAG^RAm/vglut2^ stimulation over the baseline inspirations. The portions of the abnormal inspiratory vocalization among the PAG^RAm/vglut2^-induced vocalizations (n=5, GFE3; n=4, GFP, bottom). No inspiratory vocalization was detected in the GFP control mice.

To validate the functional relevance of the anatomical connections identified above, we decided to block inhibitory inputs to RAm^VOC^ neurons. Based on the circuit diagram, we predicted that disinhibited RAm^VOC^ would provide stronger and tonic excitatory drive to MN^VOC^, that counters the inhibitory drive from the preBötC, such that vocal cord adduction may happen even during inspiration. Furthermore, if the activity of disinhibited RAm^VOC^ was sufficiently elevated, spontaneous vocalization (in the absence of social interactions) might occur. To this end, we expressed GFE3 in glutamatergic RAm^VOC^ neurons using CANE (RAm^VOC^-GFE3 mice), with RAm^VOC^-GFP mice as control (Fig. 5D). This was achieved by injection Cre-dependent CANE-hSyn-DIO-tTA together with AAV-TRE3G-GFE3 (or GFP) in the RAm in TVA/Vglut2-cre male mice after bouts of courtship USVs. GFE3 is a ubiquitin ligase specifically targeting the inhibitory post-synaptic scaffolding protein gephyrin for degradation (*37*), thereby reducing phasic synaptic inhibition onto RAm^VOC^ neurons. A previous study showed that optogenetic stimulation of RAm-projecting PAG neurons (PAG^RAm^) could reliably elicit USVs in awake head-fixed mice (*5*), a preparation allowing concurrent measurement of vocalization and breathing. Therefore, in the same RAm^VOC^-GFE3 or control mice, we also expressed ChRmine in RAm-projecting *Vglut2^+^* PAG neurons (PAG^RAm/vglut2^) using an intersectional strategy (Fig. 5D). In control RAm^VOC^-GFP mice, continuous pulses of opto-stimulation of PAG^RAm/vglut2^ reliably elicited USVs only during post-Ins, and the post-Ins was periodically interrupted by inspiration flows (Fig. 5E-F upper panels). In addition, the peak flow values for the inspiration increased during the optogenetic PAG stimulation (123.1±6.1%, n=4 mice, Fig. 5E and G), suggesting PAG^RAm/vglut2^ activation enhances inspiration (likely for inhaling sufficient air for vocalization). By contrast, in RAm^VOC^-GFE3 mice, the inspiratory interruption of vocalization was severely compromised during PAG^RAm/vglut2^ activation (Fig. 5E, lower panels). Notably, the amplitude of the few intervening inspiration during PAG stimulations was significantly reduced compared to the average inspiration peak before stimulation (49.6±10.5%, n=5 mice, p=0.020, Mann-Whitney U test for GFE3 vs GFP mice, Fig. 5F and G, lower panels). In addition, we observed that asthma-like vocal sounds were produced during the inspiration periods (21.8±5.4%, n=5 mice, Fig. 5F, gray-shaded region, and 5G), while these abnormal inspiratory vocal sounds were never observed in the RAm^VOC^-GFP control mice during PAG^RAm/vglut2^ activation. Thus, removing inhibitory synaptic inputs to RAm^VOC^ neurons compromises inspiration-vocalization coordination. The reduced inspiration amplitude is likely caused by persistent vocal cord adduction, due to a tonic excitatory drive from the disinhibited RAm^VOC^. This persistent vocal cord adduction during inspiration could also explain the abnormal asthma-like inspiratory vocalizations. Finally, consistent with the idea that tonic activation of disinhibited RAm^VOC^ neurons would cause spontaneous vocal cord closures, RAm^VOC^-GFE3 mice also produced occasional spontaneous USVs in the absence of social contexts (0.50±0.15 VOC/s, n =6 mice, fig. S4), whereas control male mice almost never utter spontaneous USVs.

## Discussion

This study reveals a vocalization-specific laryngeal premotor population in the RAm region of the caudal hindbrain and the associated neural mechanisms that coordinate vocalization with ongoing breathing. It was previously debated whether the neural circuits for laryngeal adduction required for vocal production are distributed across the ventral brainstem (*10*) or localized in one small area, such as the RAm (*38*). Considering the two results: complete mutism in all frequency ranges from RAm^VOC^ inhibition and the species-specific USV production through optogenetic activation of RAm^VOC^, we propose that RAm^VOC^ neurons form the critical laryngeal premotor node for vocal cord closure and vocalization.

Note that male USVs during courtship behaviors were used for Fos-expression to target RAm^VOC^ neurons with the CANE method. Optogenetic stimulation of RAm^VOC^ neurons was sufficient to produce and only produced USVs in male mice. Surprisingly, inhibition of the RAm^VOC^ neurons not only abolished USVs in social contexts but also audible squeaks during aversive states (tail-pinch or foot-shock). USVs and audible-squeaks in rodents have different acoustic features. USVs lie above ultra-sonic range (> 20kHz) with pure tones (*39, 40*), and rodents use aerodynamic mechanisms to produce USVs (*33–35*), while audible-squeaks occupy a human hearing frequency range (below 20kHz), with harmonics (*41*). Therefore, our finding suggests that both types of sound production in mice require the RAm^VOC^ premotor populations for laryngeal adduction, although perhaps to a different extent. Squeaks likely require additional circuit elements, such as those driving strong air exhalation, which are not activated or recruited by RAm^VOC^.

Since vocalization engages the larynx, whose opening is necessary for ongoing respiration, particularly for inspiration, laryngeal closure for sound production needs to be precisely controlled and coordinated with respiration. We show that RAm^VOC^ activation incorporates USV syllables into post-Ins, instead of violating vocal-respiration coordination. Longer USV syllables were accompanied by extended post-Ins during prolonged RAm^VOC^ activation, and the extended post-Ins was periodically interrupted by full inspiration peaks. However, it is yet to be determined how the brain volitionally prolongs RAm^VOC^ activity for syllable duration control, given that the PAG activation does not alter syllable length. It will be interesting to know whether and how the other recently identified brainstem vocal modulatory loci, such as the iRO in neonate mice (*12*) and the vocalization-related parvicellular reticular formation (VoPaRt) in rats (*14*) interact with RAm^VOC^ to control syllable patterns and durations.

Failure to coordinate laryngeal adduction and ongoing respiration could lead to vocal cord dysfunction (*42, 43*). In our proposed neural circuit for vocal-respiration coordination, inspiration-related inhibitory inputs to RAm^VOC^ can override RAm^VOC^-mediated vocal cord closure and stop ongoing vocalization. The phenotypes of disinhibited RAm^VOC^ in RAm^VOC^-GFE3 experiments, i.e., reduced inspiration peaks, hoarse sound production in inspiration phases, and spontaneous vocalization in male mice, all corroborate our proposed mechanism on vocal-respiration coordination. Our study suggests that malfunction of inhibition in the brainstem might be one type of neurological origins contributing to vocal cord dysfunction.

## Acknowledgments

We thank David Kleinfeld, Vincent Prevosto, Paul Thompson for critically reading the manuscript. We thank the Wang lab members for many technical help and suggestions during the entire process of this research project. We thank Richard Mooney lab for stimulating discussion on vocalization circuits and mechanisms.

## Author contributions

F.W. and J.P conceptualized the project, designed experiments, and wrote the paper with input from all authors. J.P. performed the majority of experiments. J.P. and J.T. analyzed data. S.C. performed histology works. S.Z., A.H., and J.T. produced key viral vectors used in this study. B.H. and S.C. provided animal husbandry. F.W. supervised all the work.

## Data and materials availability

All raw data described in this study are available from the corresponding authors upon request.

## Supplementary Materials

Materials and Methods

Figs. S1 to S4

Movies S1 to S3

References

---

**Fig. S1.**
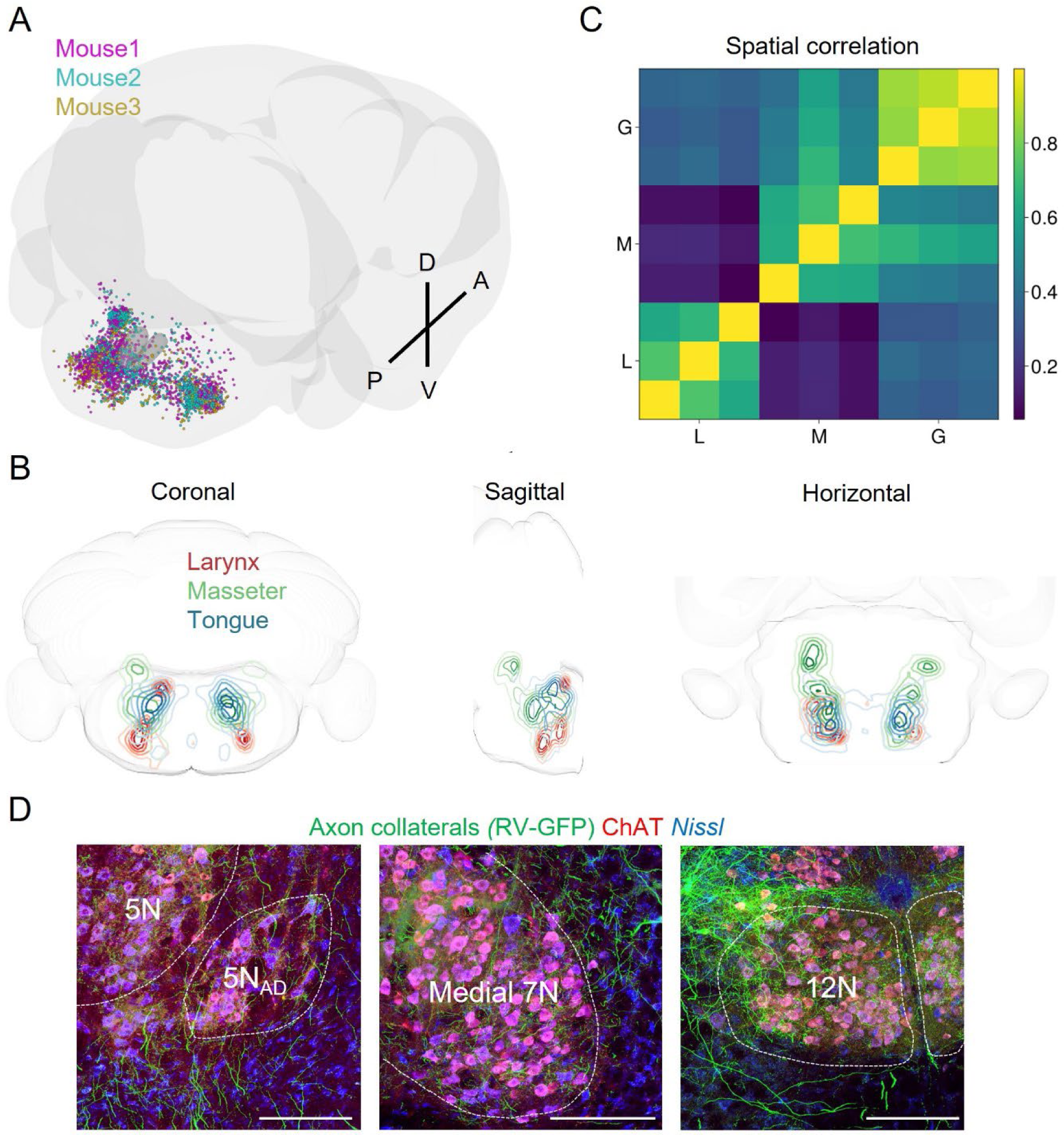
Distribution of laryngeal premotor neurons and their collaterals to the other orofacial motor nuclei. (A) 3D reconstructed distribution of laryngeal premotor neurons (magenta, aqua, and gold for 3 different mice) reconstructed in the CCF. (B) Spatial density distributions of the laryngeal premotor neurons (red) together with masseter (green) and genioglossus (blue) premotor neurons. (C) Cross-correlation of 3D spatial distributions of three orofacial premotor neurons (three examples for each: L (Larynx), M (Masseter), and G (Genioglossus). (E) Axon collaterals of rabies-labeled laryngeal premotor neurons (green) in other orofacial motor nuclei. The motor nuclei were revealed with ChAT immunolabeling (red). Neurotrace Blue was used to visualize neuronal structures. All scale bars indicate 200 µm.

**Fig. S2.**
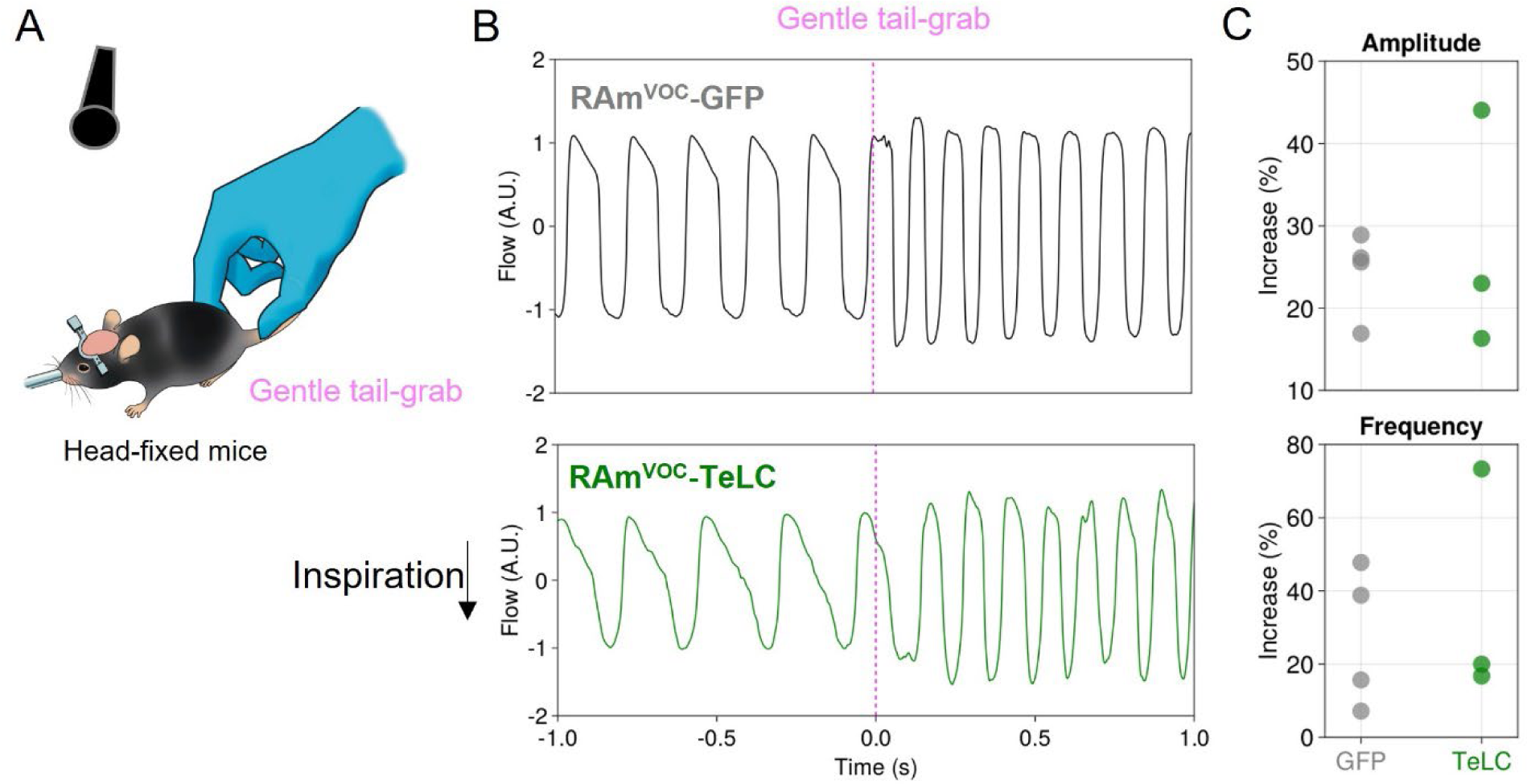
RAm^VOC^-TeLC mice were still able to modulate respiratory activity during running. (A) A scheme for encouraging mice to run for respiratory modulation. (B) Respiratory recordings in RAm^VOC^-GFP (upper) and RAm^VOC^-TeLC (bottom) mice. Magenta lines indicate the onsets of tail-grabbing. Note that downward flows correspond to inspiration. (C) Changes in amplitude (upper) and frequency (bottom) of respiratory flows. RAm^VOC^-TeLC mice (green, n=3) and RAm^VOC^-GFP control mice (grey, n=4).

**Fig. S3.**
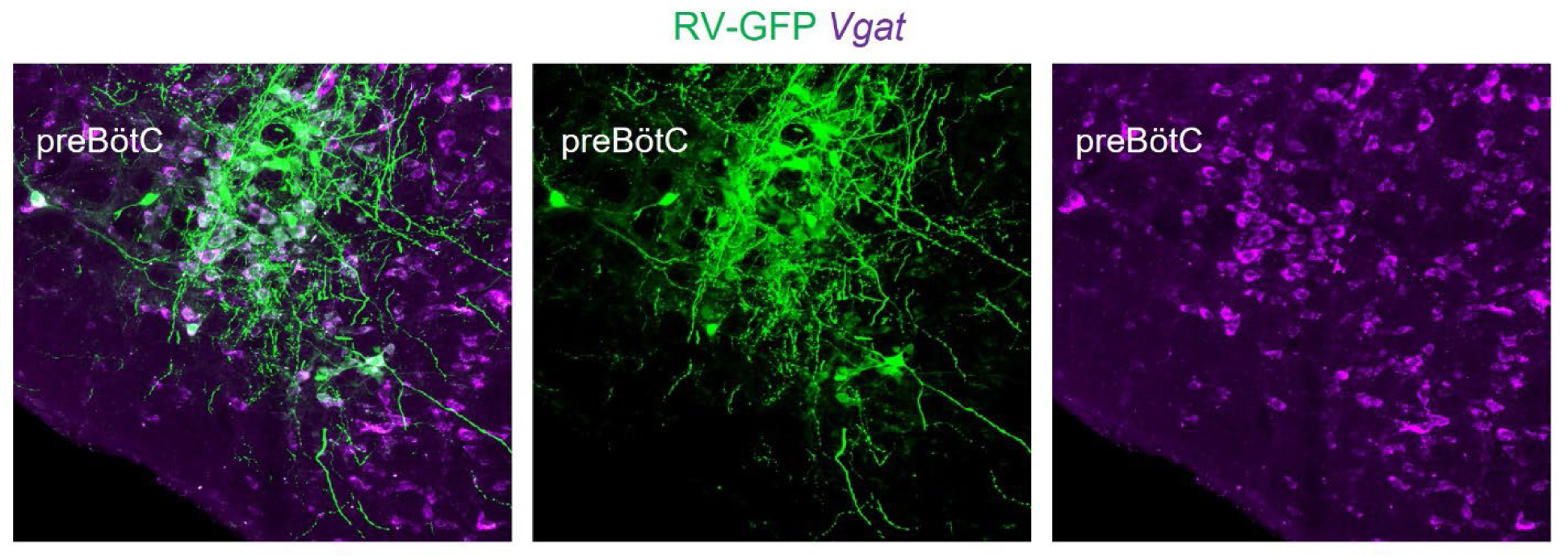
preBötC contains *Vgat*+ inhibitory laryngeal premotor neurons. Green neurons are laryngeal premotor neurons labeled with monosynaptic rabies-GFP through transsynaptic tracing (as described in Figure 1), and the in-situ hybridization signal with *Vgat* probe is shown in magenta.

**Fig. S4.**
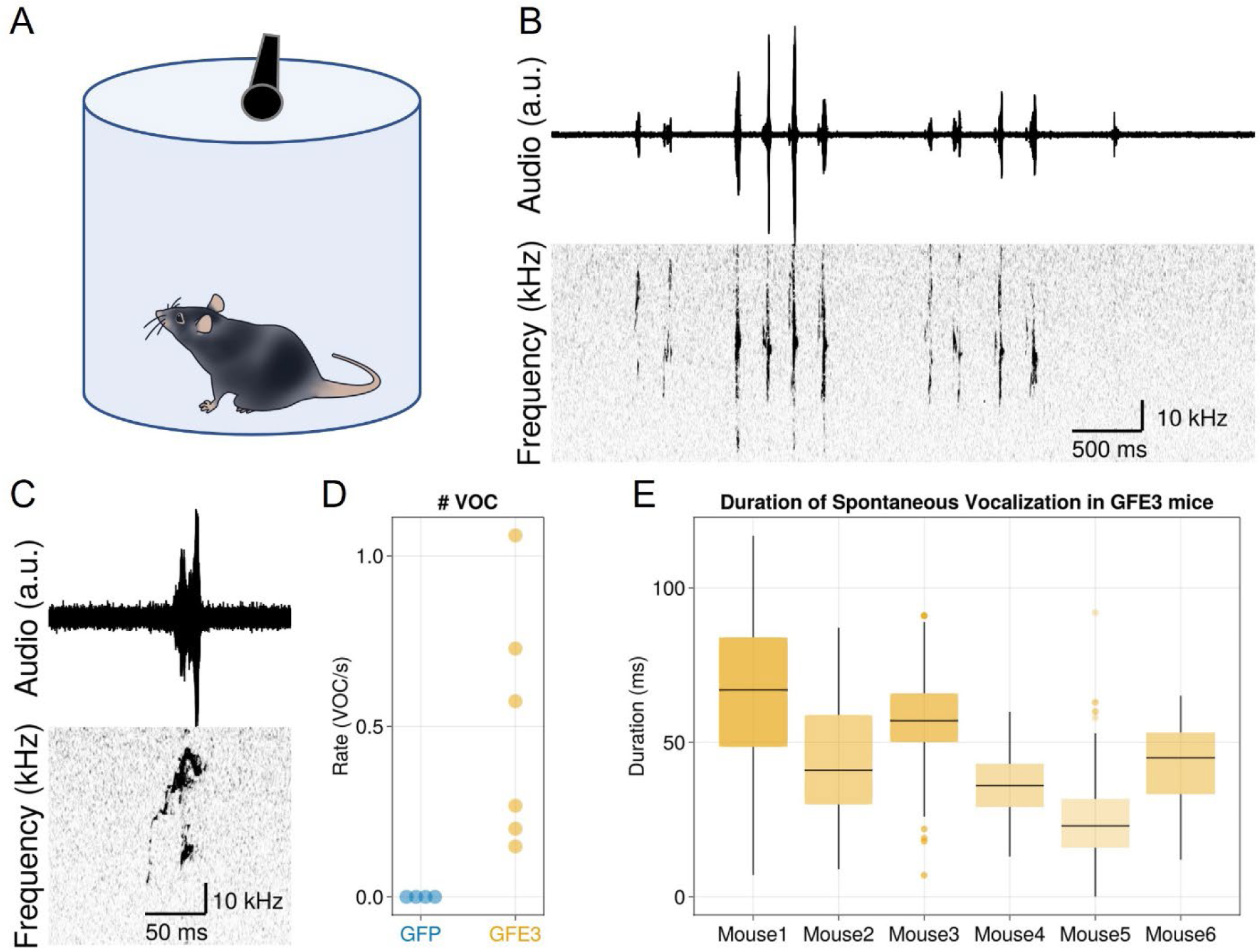
Spontaneous vocalizations emerged in the RAm^VOC^-GFE3 mice. (A) Schematic for recording male mice vocalization in an isolation chamber. (B) Raw audio data (upper) and corresponding spectrogram plots (bottom). (C) An example of a single USV syllable from the spontaneous vocalizations. (D) Rates of spontaneous vocalizations in RAm^VOC^-GFE3 mice (n=6) and RAm^VOC^-GFP mice (n=4). Note that no spontaneous vocalization was found in the GFP control mice. (E) Durations of the spontaneous vocalizations of six RAm^VOC^-GFE3 mice. Single dots show outliers.

**Movie S1. A RAm^VOC^-GFP mouse in a foot-shock chamber.** Raw audio signals (upper left) and corresponding spectrogram (bottom left, in ultrasonic range). Sounds in the video (right) were recorded with an audible microphone.

**Movie S2. A RAm^VOC^-TeLC mouse in a foot-shock chamber.** The mouse showed behavioral responses to foot-shock but did not produce audible squeaks.

**Movie S3. Glottal responses to RAm^VOC^-ChRmine stimulation.** Four red dots are used to trace the glottal area. Red lines represent the glottal area. Green bars indicate the optogenetic stimulation periods.

